# The interplay between demography and neutral evolution at the expansion front of a widespread conifer, *Picea sitchensis*

**DOI:** 10.1101/327742

**Authors:** Joane S. Elleouet, Sally N. Aitken

**Affiliations:** Department of Forest and Conservation Sciences, 2424 Main Mall, University of British Columbia, Vancouver, BC, V6T1Z4, Canada

## Abstract

Tree species in the northern hemisphere have advanced and retreated with interglacial and glacial periods, and are currently subject to rapid anthropogenic climate change. These observations prompt questions about the mechanisms allowing tree populations to respond quickly to selection pressures when establishing into new areas. Focusing on the northern expanding range edge of *Picea sitchensis*, a widespread conifer of western North America, we ask how genetic structure and diversity develop during colonization, and assess the role of demographic history in shaping the evolutionary trajectory of a colonizing population. By combining nearly 500 years of tree-ring and genetic data at the expansion front on the Kodiak Archipelago, we show that allelic richness - but not expected heterozygosity - increased rapidly during early stages of establishment in the 1600s, while genetic differentiation from populations further from the front decreased. This trend ended in the 1700s, after an increase in population growth rate. These findings highlight the major role of long-distance pollen dispersal in the recovery of genetic diversity during initial stages of colonization, and suggest that demographic dynamics including an initial lag in population growth are likely limiting factors in the adaptation of tree populations tracking their niche in a changing climate.

## Introduction

With an enhanced understanding of plant species migrations during past postglacial cycles and numerous observations of current climate change effects on species distributions, we are recognizing more than ever before the ubiquitous nature of range shifts. This awareness comes with a substantial literature including simulation studies of genetic changes during range expansions (Bialozyt et al., 2006; Hallatschek & Nelson, 2010; Peischl et al., 2013), empirical studies of expanding species and associated evolutionary patterns (e.g. Darling et al., 2008; Pujol & Pannell, 2008), and metaanalyses and reviews highlighting the general patterns common to - or variable among - phylogenetic and functional biological groups (Excoffier et al., 2009). Successive founder effects along colonization routes are a well-studied phenomenon causing an erosion of diversity and enhancing genetic differentiation (Excoffier et al., 2009; Hewitt, 2000; Slatkin & Excoffier, 2012). However, mechanisms related to species’ dispersal and life history traits have been shown to influence the genetic outcome of range expansions and give rise to fundamentally different spatial patterns of genetic structure (Bialozyt et al., 2006; Waters et al., 2013). It is therefore not surprising that among organisms with different life history and dispersal traits, genetic effects of range expansions are not consistent (Eckert et al., 2008).

This heterogeneity is present across studies of tree species. Empirical studies at the scale of species ranges have found both higher genetic diversity (Born et al., 2008; Pluess, 2011; Shi & Chen, 2012) and lower genetic diversity (Johnson et al., 2017; Kitamura et al., 2015; Marsico et al., 2009; Mimura & Aitken, 2007) in leading edge populations of tree species after range expansion. Temperate forest tree species are generally associated with high gene flow via wind-borne pollen across large geographic distances (Kremer et al., 2012), as well as a long lifespan and juvenile phase. These characteristics can have a strong influence on the interplay between genetic and demographic processes during range expansion. Founder effects may be buffered by high levels of gene flow (Austerlitz et al., 1997). The longevity of trees allows for founders to persist until other propagules colonize, and their long juvenile phase forces the reliance on gene flow via foreign pollen, providing genetic diversity to the establishing population (Austerlitz et al., 2000). In addition, long-distance dispersal can prevent the erosion of genetic diversity along expansion routes (Le Corre & Kremer, 1998), promote genetic differentiation between demes (Austerlitz & Garnier-Géré, 2003) or suppress local introgression (Amorim et al., 2017).

Recent empirical studies involving exhaustive sampling and pedigree reconstruction in disjunct forest stands at range edges have greatly enhanced our understanding of demographic mechanisms shaping the genetic composition of expanding populations at the local scale. Founding individuals, often arriving via long distance dispersal of seeds, play a major role at the start of establishment. Troupin et al. (2006) found that spatial genetic structure of a population shortly after range expansion strongly reflected the genetics of founding trees in a *Pinus halepensis* population. Lesser et al. (2013) identified Allee effects in a new *Pinus ponderosa* stand, highlighting the importance of a certain level of long-distance dispersal of founders in the establishment success of the new population. A general finding in these studies is the predominance of high levels of pollen flow shortly after founding, leading to a quick recovery of genetic diversity during recruitment (Hampe et al., 2013; Lesser et al., 2013; Pluess, 2011; Sezen et al., 2007).

Here, we take a multi-scale approach to the study of expanding tree populations, combining observations of spatial and temporal patterns at the regional and local scale. In particular, we ask how demographic and spatial patterns of colonization affect genetic structure along an expansion route in space and time, and interact with potential geographic barriers to gene flow. To do so, we combine nearly five centuries worth of demographic and genetic data from several forested sites along the recent colonization route of a widespread North-American conifer, Sitka spruce (*Picea sitchensis* (Bong.) Carr.). With its ability to withstand salt spray and thrive in hypermaritime environments, *P. sitchensis* is a major component of Pacific coastal forests. Although it is rarely the most abundant species in the southern and central part of its range, it dominates the forest cover together with mountain hemlock on the Kenai Peninsula and is the only forest tree species on the Kodiak Archipelago. Historical human records report a rapid advance of the monospecific *P. sitchensis* forest on the Kodiak Archipelago (Griggs, 1914; Vincent, 1964). The westernmost and largest island of the group, Kodiak Island, is thought to have been colonized no more than 500 years ago (Griggs, 1937). *P. sitchensis* is found in dense stands on Afognak Island and the forest density tapers south-westward towards small groves of young trees (Tae, 1997). The absence of spruce pollen in paleoecological records on Kodiak Island (Bowman, 1934) strongly suggest that this is the first occurrence of the *P. sitchensis* forest at this site since the last glacial period. This range expansion therefore seems to be part of the long-term post-glacial colonization process of the species, with the most recent front advance having likely been facilitated by a nearby volcanic eruption in 1912, which reduced interspecific competition between *P. sitchensis* seedlings and herbaceous species (Tae, 1997). Southwestern Kodiak Island features tundra grasses and scattered shrubby forms of *Alnus viridis, Populus trichocarpa*, and *Betula nana.* Earlier studies focusing on *P. sitchensis* detected asymmetric gene flow from core Alaskan populations to the Kodiak Archipelago (Holliday et al., 2012), as well as a high self-fertilization rates (Gapare & Aitken, 2005; Mimura & Aitken, 2007) and a lack of adaptive potential (Lobo, 2011) on Kodiak Island. Based on this knowledge, we propose to characterize gene flow from populations on the Kenai Peninsula to the Kodiak Archipelago and identify demographic or genetic mechanisms responsible for reduced levels of genetic diversity at the expansion front.

Using dendrochronological methods, we first conduct a demographic analysis over island and continental regions to determine the timing and spatial structure of dispersal patterns during range expansion on the Kodiak Archipelago. To assess the current and past extent of differentiation and genetic diversity at the regional level, we then quantify genetic population structure between regions and genetic diversity within regions for different age classes at the northern range of *P. sitchensis*. This direct monitoring of genetic diversity and structure allows for the quantification of the extent and duration of potential founder effects, as well as the relative importance of early colonizers and subsequent gene flow in the accumulation of genetic diversity of the growing population. Finally, we take a closer look at sites at the expansion front, to determine the short-term genetic consequences of fine-scale dispersal patterns and demography.

## Materials and Methods

### Sampling design

We focused on the northern range of *P. sitchensis* in south-central Alaska. In 2013 and 2015, fifteen sampling sites in healthy forests with old-growth characteristics and with no evidence of past outbreaks of spruce beetle (*Dendroctonus rufipennis*) were sampled on Kodiak Island, Afognak Island, and on the Kenai Peninsula near Seward (Figure 1a). Sample sizes within sites varied between 12 and 86 trees, depending on the size of the sampled site and its accessibility (Table 1). We classified trees into four canopy structure levels (large canopy trees, medium-sized canopy trees, sub-canopy trees, and immature saplings) and sampled sites to maximize the range of tree ages and to obtain even sample sizes from each canopy structure level. Large canopy trees were typically >70cm in diameter, and showed growth forms consistent with earlier growth in an open environment (numerous large dead branches low on the trunk and strongly tapered stems), especially on the Kodiak Archipelago. Medium-sized canopy trees were generally <70cm in diameter and showed no signs of open growth. Sub-canopy trees were mature trees that had not reached the main canopy, and immature saplings were generally no more than 2m tall. All sampled trees were separated by at least 150 meters to avoid high relatedness between individual samples. For DNA extraction, we sampled young needles whenever possible; when foliage was out of the reach of a pole pruner we took two 1 cm-diameter cambium disks with a leather punch. Sampled materials were stored in paper envelopes in silica gel until DNA extraction. To estimate the age of individuals sampled, we used an increment borer to core each tree up to 5 times as low as possible to obtain a wood sample that included the pith or signature thereof. A detailed description of dendrochronological methods is available in the supplemental information. The age of saplings was approximated by counting major branch clusters on the stem. We used available samples of needles from 15 canopy trees in two additional sites on Shuyak Island and Port Chatham for inclusion in population genetics analyses but did not have tree ring data for these populations.

**Figure 1.**
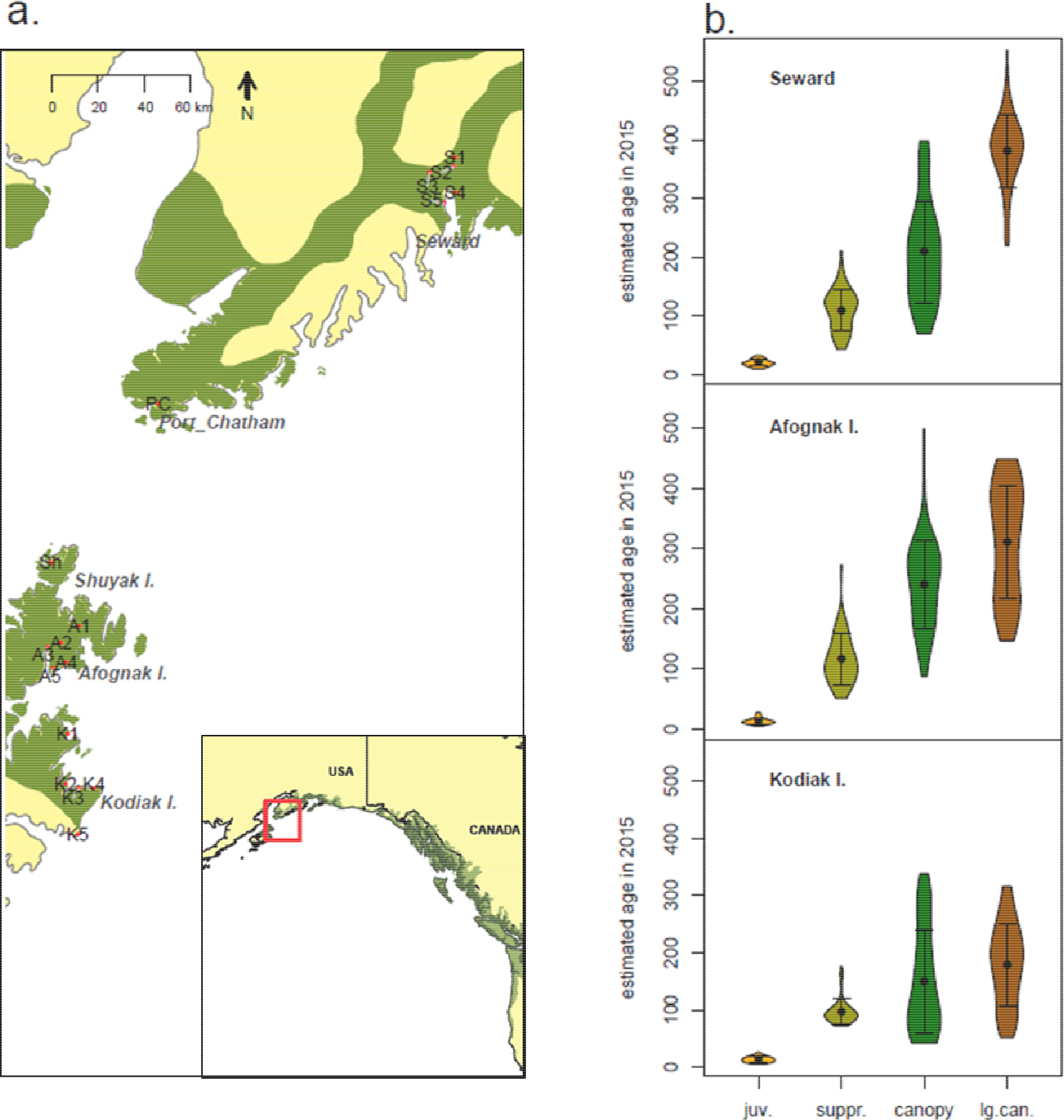
**a.** Sampled sites in south-central Alaska, with the whole range of *P. sitchensis* (green area) in inset. **b.** Violin plots of individual tree ages for each canopy structure level within each region sampled, with mean and standard deviation displayed in black.

**Table 1.** Nomenclature, description, and diversity estimates of sampled sites (shown on Figure 1a). Size=sample size; H_e_=population-level heterozygosity averaged across loci; mean canopy age=average tree age across large canopy and medium-sized canopy trees, in years.

| site | region | latitude | longitude | size | $H_e$ | mean canopy age |
| --- | --- | --- | --- | --- | --- | --- |
| S1 | Seward | 60.253 | -149.357 | 37 | 0.283 | 294 |
| S2 | Seward | 60.213 | -149.369 | 36 | 0.29 | 288 |
| S3 | Seward | 60.19 | -149.558 | 48 | 0.268 | 285 |
| S4 | Seward | 60.107 | -149.354 | 48 | 0.251 | 319 |
| S5 | Seward | 60.06 | -149.444 | 26 | 0.202 | 266 |
| PC | Port Chatham | 59.224 | -151.703 | 15 | 0.123 | NA |
| Sh | Shuyak | 58.563 | -152.555 | 15 | 0.183 | NA |
| A1 | Afognak | 58.298 | -152.329 | 24 | 0.119 | 206 |
| A2 | Afognak | 58.231 | -152.475 | 37 | 0.155 | 223 |
| A3 | Afognak | 58.209 | -152.585 | 12 | 0.12 | 315 |
| A4 | Afognak | 58.145 | -152.433 | 25 | 0.121 | 239 |
| A5 | Afognak | 58.118 | -152.541 | 86 | 0.15 | 310 |
| K1 | Kodiak | 57.845 | -152.422 | 52 | 0.134 | 203 |
| K2 | Kodiak | 57.638 | -152.437 | 49 | 0.134 | 147 |
| K3 | Kodiak | 57.619 | -152.334 | 50 | 0.123 | 207 |
| K4 | Kodiak | 57.617 | -152.219 | 31 | 0.136 | 201 |
| K5 | Kodiak | 57.428 | -152.344 | 48 | 0.13 | 66 |

### Dating population establishment

We used two methods to estimate timing of population establishment at the site level: canopy age and date of canopy closure. Canopy age was obtained by averaging ages of all canopy trees based on tree ring counts. This estimate has the advantage of being accurate for canopy age but does not necessarily reflect initial forest establishment if tree longevity is shorter than the duration of established forests since colonization or between major stand-replacing disturbances. Canopy closure time is an attempt to overcome this limitation and relies on signatures of growth during early life stages, based on the hypothesis that individuals growing in an open environment will experience faster growth in early stages of life compared to individuals competing under a closed canopy. Therefore we expect: (1) that trees on sites that were forested long before the time encompassed by our sampling will have relatively narrow growth rings for all samples at the juvenile stage; (2) that sites still in the process of forest colonization will show wide growth rings for most samples at the juvenile stage; (3) that formerly forest-free sites will show a shift from wide to narrow growth rings in juveniles during colonization as the canopy closes.

For each tree core, we measured and averaged annual growth increment between years 10 and 20 (to avoid signals of competition with small-statured shrubby or herbaceous plants). We modeled the relationship between individual estimated establishment date *x* and individual tree average annual juvenile growth *y* at the site level using two different regression models: a linear model *y~ax* + *b* and a logistic model of the type *y~ a*/(1 + *e*−^*b*(*x*−*h*)^. We expect sites following cases (1) and (2) above to show a better fit with a linear model with insignificant or weak slope, and sites following case (3) to show a better fit with the logistic model. We retained the model with the lowest AIC. When sites were best fitted by a logistic model, we checked that the relationship between year of establishment and growth increment was negative (*b*<0) and recorded *h* as the estimated date of canopy closure. When the linear model was the best fit, we tested the significance of the slope (*p*-value for coefficient *b*). Site K4 was removed from this analysis due to the sub-canopy structure level being under-represented.

### Genotyping

To obtain putatively neutral markers for 639 trees, we used genotyping-by-sequencing (GBS) with a sbf1-msp1 double-digest protocol (Elshire et al., 2011). Libraries were sequenced with the HiSeq 2000 system, producing 100-bp single reads. We aligned the filtered reads to the white spruce reference genome WS77111_V1 (Warren et al., 2015) using the bwa mem alignment algorithm. Alignment files were then input into a variant calling pipeline using functions from the program GATK (McKenna et al., 2010). A more detailed description of the bioinformatics processing steps is provided in the supplementary materials. After SNP calling, we removed all singletons across the 639 genotyped diploid individuals. Finally, when several SNPs were present less than 100bp apart, we retained only one of them, resulting in a final dataset with genotypes for 3244 biallelic SNPs. Different subsets of SNPs were used for different population genetics analyses, depending on selected individuals and chosen missing data threshold. Although the GBS approach outputs datasets with considerable missing data, it also provides a cost-effective genome-wide picture of genetic diversity and structure.

### Visualizing population structure

We visualized population structure among all regions sampled using 2 methods: principal component analysis (PCA) and STRUCTURE clustering (Falush et al., 2003; Pritchard et al., 2000). For the PCA, we retained all genotype calls and filtered sites for missing data with a 40% cutoff value. The resulting dataset included 639 *P. sitchensis* individuals genotyped for 220 SNPs. We replaced all missing data by their mean over all individuals prior to PCA. The R packages *adegenet* and *ade4* were used to convert data files and perform a centered PCA. To characterize further population structure within the dataset, we used the program STRUCTURE, which detects clusters of individuals based on Hardy-Weinberg equilibrium within clusters. We first performed exploratory runs with run lengths from 10k to 100k after a 10k burnin and 3 replicate runs for each run length. For K≤4, a run length of 50k was sufficient, whereas for K>4, a run length of 100k was necessary. We used the independent allelic frequency model and the admixture ancestry model for all runs, and performed 3 runs for each value of K between 1 and 6.

### Temporal patterns of diversity and structure

To infer the role of founder individuals and subsequent migration in the development of genetic diversity of the Kodiak-Afognak population, we selected all sites with less than 20% missing genotypic data for trees with age estimates. This dataset (120 SNPs, 412 trees) was used to estimate the year of first observation of each allele in the growing population. A 1000-replicate randomization was applied to model the random distribution of allele accumulation curves against which to test significance of the observed results. To reconstruct the changes in gene flow patterns between continental and island populations, we assessed pairwise population differentiation between regions by computing the Weir and Cockerham (WC) F_ST_ estimator with R functions modified from the *adegenet* and *hierfstat* packages. Loci with less than 30% missing data over the whole sample were used. As the WC estimator is sensitive to unbalanced sample sizes (Bhatia et al., 2013), we randomly subsampled the larger population sample to the size of the smallest population sample. Confidence intervals around F_ST_ estimates were assessed with 1000 bootstraps.

### Site-level summary statistics

Using SNPs that were well represented in each region (< 60% missing data across the sample), we calculated expected heterozygosity, F_ST_ and allelic richness at the site level using the R packages *adegenet, hierfstat* and *PopGenReport*, respectively. The latter uses the methods of El Mousadik and Petit (1996), which corrects for variable sample sizes through rarefaction. Changes in dissimilarity between sites over time on the Kodiak Archipelago were estimated using the dissimilarity calculations of Petkova et al. (2016). Estimates at the individual tree level rather than allele frequencies (such as F_ST_) better suited our temporal analysis at the local scale due to low within-site sample sizes for each cohort. Briefly, we computed a matrix of genetic distance between pairs of individuals using the average squared genetic difference across all well-represented SNPs (<50% missing data). We then calculated D, the mean genetic distance over all possible pairs of individuals from 2 distinct sites, as a measure of pairwise dissimilarity between sites. To avoid confounding effects of within-site differences and better represent genetic differences resulting from gene flow variations, we calculated between-site dissimilarity (D_b_) by subtracting the average within-site dissimilarity from D.

## Results

### Demographic patterns from tree rings

We successfully estimated tree ages for a total of 607 samples (N=412 on the Kodiak Archipelago, N=195 in the Seward region on the Kenai Peninsula), evenly distributed among four canopy levels. Estimated tree ages ranged from 5 to 552 years. Canopy trees were generally younger on Kodiak Island (145 y.o. on average) than on Afognak Island and Seward (>200 y.o.) and ages of large canopy trees differ considerably between regions: the largest canopy trees are younger in regions closer to the range limit (Figure 1b). To obtain a finer resolution of the spatial demographic patterns, we calculated the mean canopy age at sites within regions (Figure S1). Canopy age at all Seward sites was around 250 to 300 years, with overlapping standard errors. From these relatively consistent estimates, we deduced that canopy age will not correlate with population establishment date on the Kodiak Archipelago beyond this time, as Seward sites were colonized more than 1000 years ago (Jones, 2008; Mann & Hamilton, 1995). Within Kodiak Island, two of the southernmost areas (K3 and K5) have no trees older than 200 and 135 years, respectively, with mean canopy ages of 140 and 67 years. K5 is in the southernmost *P. sitchensis* forest, located on the Southeast coast of Kodiak Island. At this site, tree ages within structure levels are highly homogenous, with most canopy trees between 30 and 80 years old. The canopy is close to the ground and there are no suppressed trees growing in the understory, suggesting that this site was recently colonized. In general, canopy age is higher on Afognak Island than on Kodiak Island. However, this pattern of decreasing canopy age towards the expansion front breaks down at the local scale: some sites in the south have an older canopy than northern sites, especially on Afognak Island (*i.e.*, A1 and A5, Figure S1). The large variability in canopy age among areas at a similar latitude (*i.e.*, K3 and K4, Figure S1) suggests that colonization on the Kodiak Archipelago occurred via patchy dispersal rather than a linear advancing wave.

We detected a signal of canopy closure using data from early-age annual growth increment patterns at the site level (Figure 2). As expected, none of the five sites on the Kenai Peninsula (S1 to S5) showed any significant relationship between growth increment and time for either the linear or the logistic model, with growth rings around 1mm wide. This is consistent with growth under a well-formed canopy with constant population density and no large-scale disturbances over the time span sampled. Contrasting with this pattern, four sites on Kodiak Island (K1, K2, K3, K5) show very large juvenile growth rings (3-7 mm) for older trees. Sites on Afognak Island show moderately large growth rings (2-4 mm) for older trees (A1 to A5). Another strikingly different pattern is that all sites but one on the Kodiak Archipelago present best fit from a logistic curve or a linear model with significant negative slope, suggesting that over time juvenile growth gradually decreased at all sites to values similar to current values in the Seward population. K5 is the only site where ring width does not decrease, with current values above 2mm. This is consistent with a young stand where all mature trees present developed in the absence of substantial intraspecific competition. For sites where a logistic curve was the best fit, estimated dates of canopy closure *h* varied little among sites. As growth rings are narrower in old trees on Afognak Island than on Kodiak Island but we still observe a decrease in radial growth over time, we suggest that stands with closed canopies on Afognak Island earlier than the period for which we have data, and that the detected signal corresponds to a slow increase in stand density.

**Figure 2.**
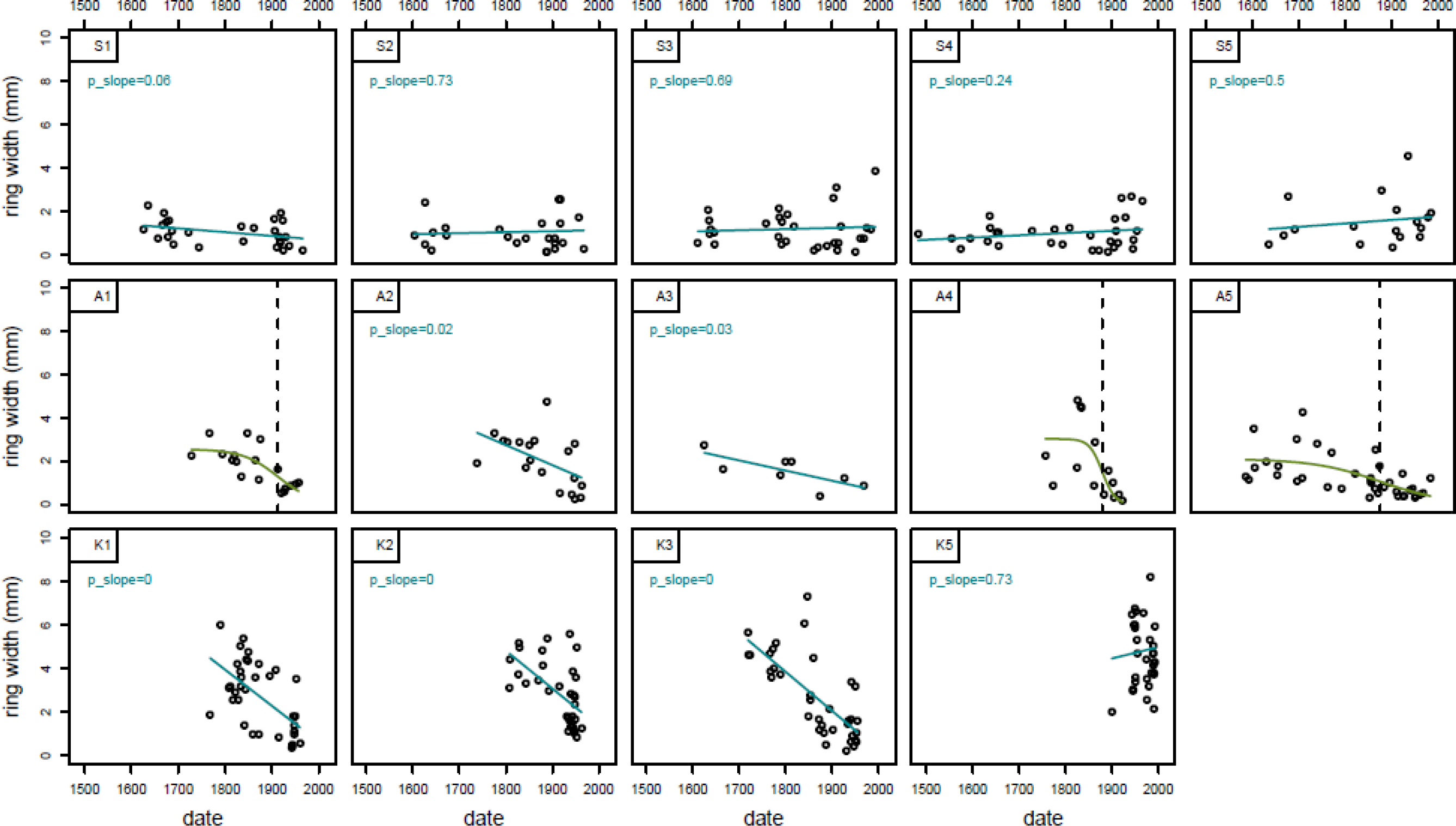
Individual annual radial growth between years 10 and 20 vs. date of establishment. The best model among linear and logistic models is displayed in blue (linear model) or green (logistic model). In the latter case the estimated time of canopy closure *h* is represented by a dashed line.

Finally, we computed the cumulative distribution of establishment dates of canopy trees in both the Kodiak Archipelago and Seward populations (Figure 3a). There was a sharp increase in the cumulative number of established canopy trees on the Kodiak Archipelago around 1700. The cumulative distribution of the Seward sample suggests that this shift is not due to intrinsic mortality in trees established before 1700. Indeed, such old trees are present in the Seward population (although few trees established before 1550 were sampled). Instead, the observed shift in age distribution on the Kodiak archipelago compared to Seward might either indicate a genuine increase in establishment rate around 1700 or an elevated extrinsic mortality of trees established before that time. However we do not know of any catastrophic climatic or geological event from this time, nor do we observe signals of it in annual tree rings of trees established before 1700 on the archipelago.

**Figure 3.**
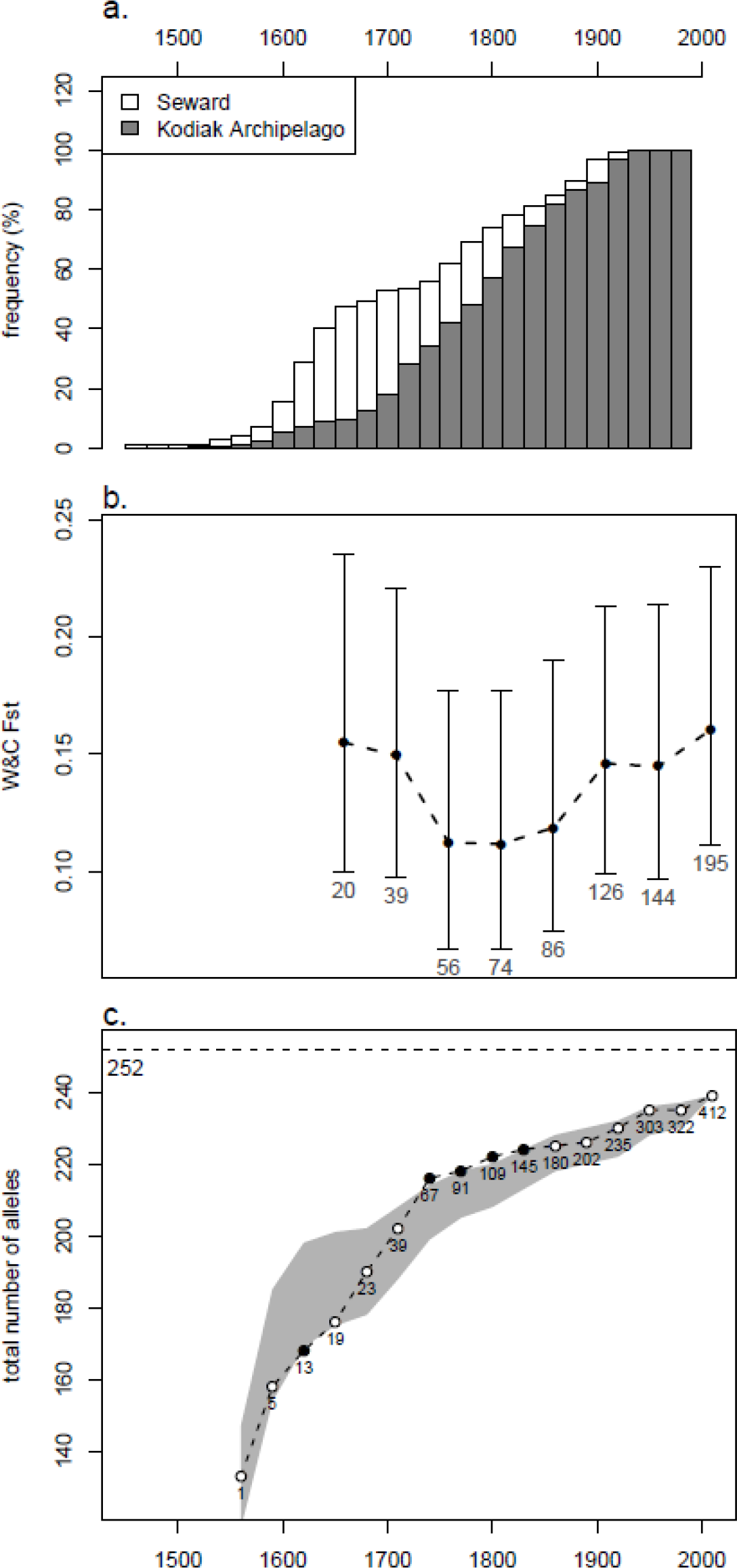
Demographic and genetic changes over time on the Kodiak Archipelago in relation to the Kenai Peninsula. **a.** Cumulative distribution of establishment dates for canopy trees for the Kodiak Archipelago and the Seward region. **b.** Temporal change in F_ST_ between the Kodiak Archipelago and Seward. F_ST_ Values are calculated for the cumulative sample (each sample associated with a date is made of all individuals alive at this date). Error bars represent confidence intervals from 1000 bootstraps. The number of individuals per population is indicated below error bars. **c.** Allele accumulation curve for the Kodiak Archipelago with 95% interval band from 1000 random permutations (grey). Numbers under datapoints correspond to cumulative sample sizes. Data points with values outside the 95% confidence band are represented with filled circles. The top dotted line correspond to the maximum number of alleles in the whole sample of 639 individuals.

### Genetic structure and diversity in space and time

Both PCA plots (Figure S2) and STRUCTURE analyses (Figure S3) suggest that population differentiation is moderate and mainly separates Seward from the other populations. In particular, the mixed ancestry of Shuyak and Port Chatham displayed in the K=2 and K=3 STRUCTURE bar plots as well as their position on the PCA plot suggest that the strait separating the archipelago from the mainland does not produce any marked differentiation pattern, at least not compared to similar overland distance.

To determine how the present regional pattern of population structure evolved, we analyzed the evolution of F_ST_ over 400 years between the Kodiak Archipelago and the Seward region (Figure 3b). Despite large confidence intervals around estimates, we observe a decrease in F_ST_ from 0.15 in 1610 to 0.12 in the mid-1700s, followed by a weak, statistically nonsignificant increase to about 0.15, the current estimate. The early decrease in differentiation could indicate relatively high gene flow from the mainland to the Kodiak Archipelago during early population establishment. A shift to local recruitment likely happened in the 1700s, putting an end to the decreasing trend in genetic differentiation. This shift is coincident with the upward shift in the distribution of establishment time of canopy trees on the Kodiak archipelago (Figure 3a).

Genetic diversity decreased towards the expansion front for both allelic richness and expected heterozygosity calculated over polymorphic loci (Figure 4). The Seward population had the highest allelic richness, and Kodiak Island, Afognak Island, and Port Chatham had the lowest. Interestingly, Shuyak Island has a higher allelic richness than Port Chatham suggesting connectivity of the Shuyak population with other populations, possibly from *P. sitchensis* forests outside of those sampled, or from *P. glauca* populations north of the archipelago. Heterozygosity is high everywhere but on Kodiak and Afognak Islands, suggesting a deficit of some alleles common elsewhere at the expansion front.

**Figure 4.**
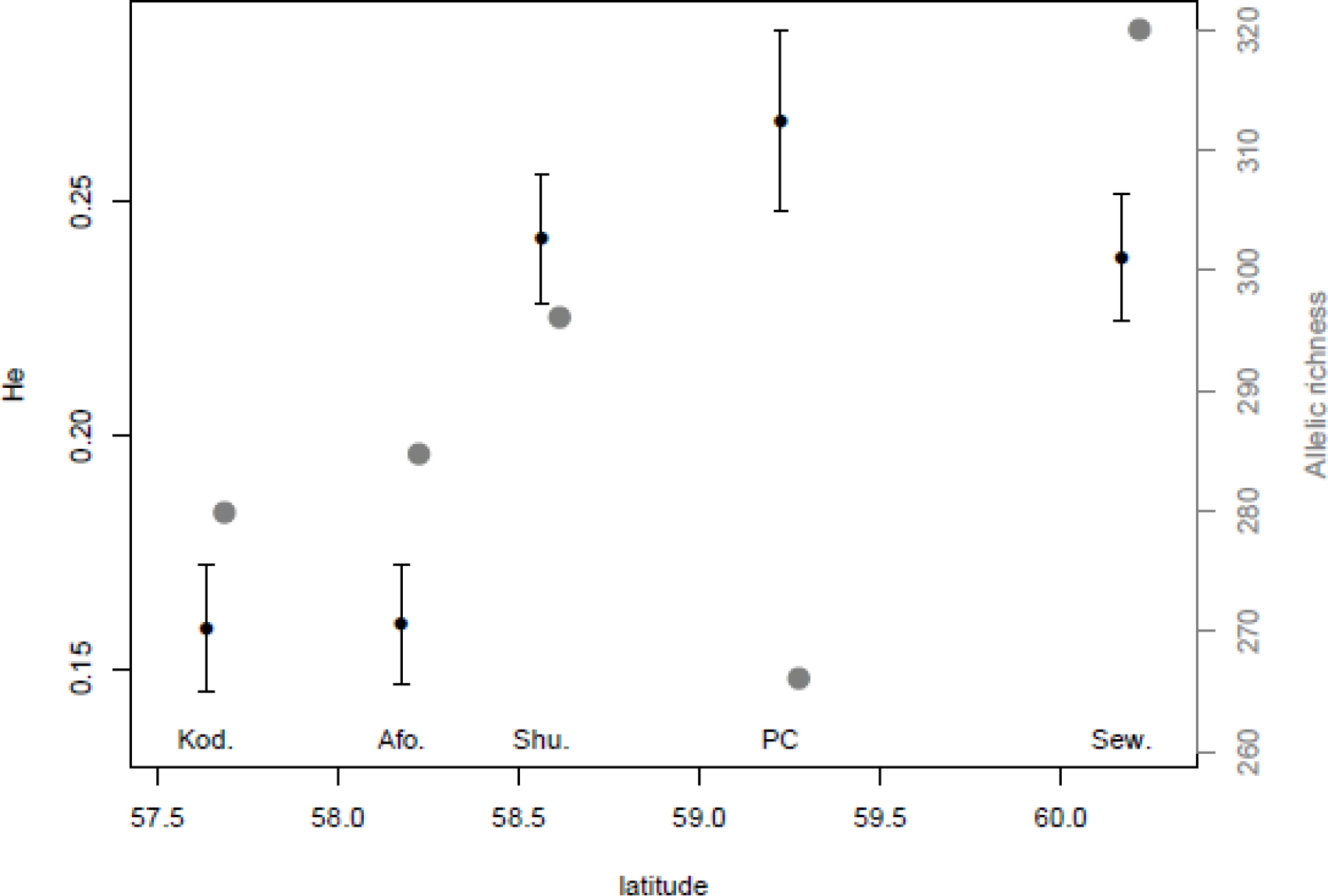
Regional expected heterozygosity over variable SNPs (black) and estimated allelic richness (grey) vs. latitude or regions. Error bars on H_e_ are standard errors of the mean. Kod.=Kodiak Island, Afo.=Afognak Island, Shu.=Shuyak Island, PC=Port Chatham, Sew.=Seward.

To determine how quickly the Kodiak-Afognak populations acquired their current allelic diversity, we built an allele-accumulation curve (Figure 3c) and compared it to a null model of comparable sample sizes. We found that most alleles present in the data were acquired between 1620 and the mid-1700s, a trend confirmed not to be an artefact of sampling effects.

### Patterns of genetic structure and diversity at the colonization front

Expected heterozygosity calculated for each sampled site using all SNPs that are polymorphic ranged from 0.11 to 0.28 across the Kodiak Archipelago (table 1). There is no evidence for a latitudinal decrease within the archipelago towards the edge of the range: we calculated a correlation coefficient of 0.04 between latitude and H_e_ (Pearson’s correlation test, *p* = 0.9004). We calculated a similar correlation coefficient value between expected heterozygosity and canopy age (*r*=0.08, Pearson’s correlation test, *p* =0.8101), and again failed to detect any erosion of genetic diversity during successive colonization of demes at the expansion front.

To test the hypothesis that colonization leads to genetic sectors on the landscape, we computed pairwise F_ST_ among sites (Figure 5). Larger F_ST_ values among areas on the Kodiak Archipelago than among areas on Seward would suggest the presence of such colonization-specific mechanisms at the front. However, we observe very low (<0.1) pairwise F_ST_ values between sites on the archipelago, similar to those observed among Seward sites. This could be due to high gene flow among sites after long-distance dispersal founding events, making genetic sectors too transient to be observable in current datasets. Alternatively, it could be due to a homogeneous pool of founding individuals across the archipelago, which could occur if all propagules came from one, already depauperate source population. To test these alternative hypotheses, we took a landscape genetics approach and calculated genetic dissimilarity among sites at different distances within the Kodiak Archipelago and at four different times, and studied the change in dissimilarity between sites over time (Figure 6). Dissimilarity between sites in 1710 was significantly higher than during subsequent centuries: We observe average dissimilarity values an order of magnitude lower in 1810, 1910 and 2010 than in 1710. This result brings support to the hypothesis that post-founding gene flow prevented initial genetic sectors from persisting.

**Figure 5.**
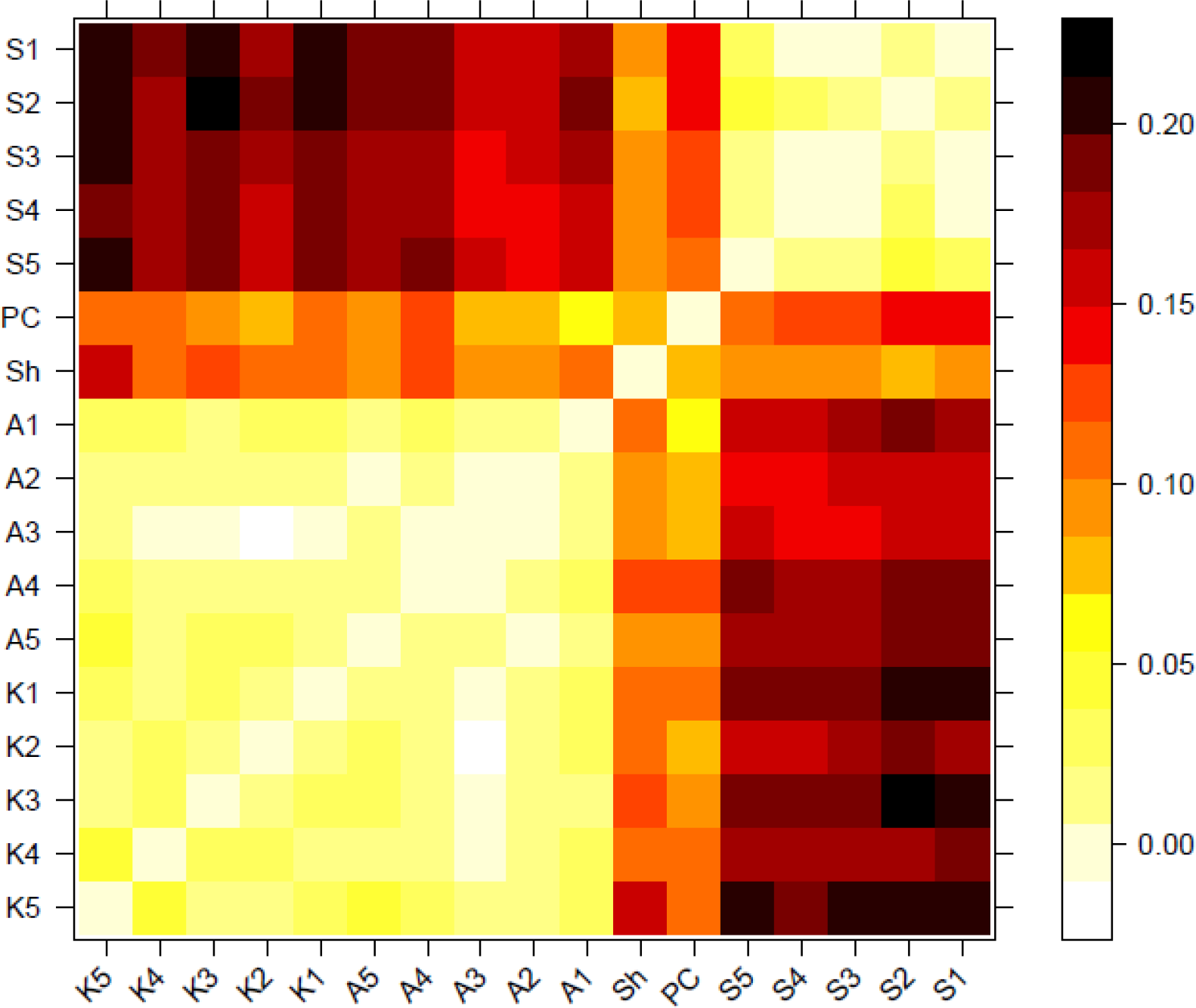
Heat map of pairwise Fst values between sampled sites.

**Figure 6.**
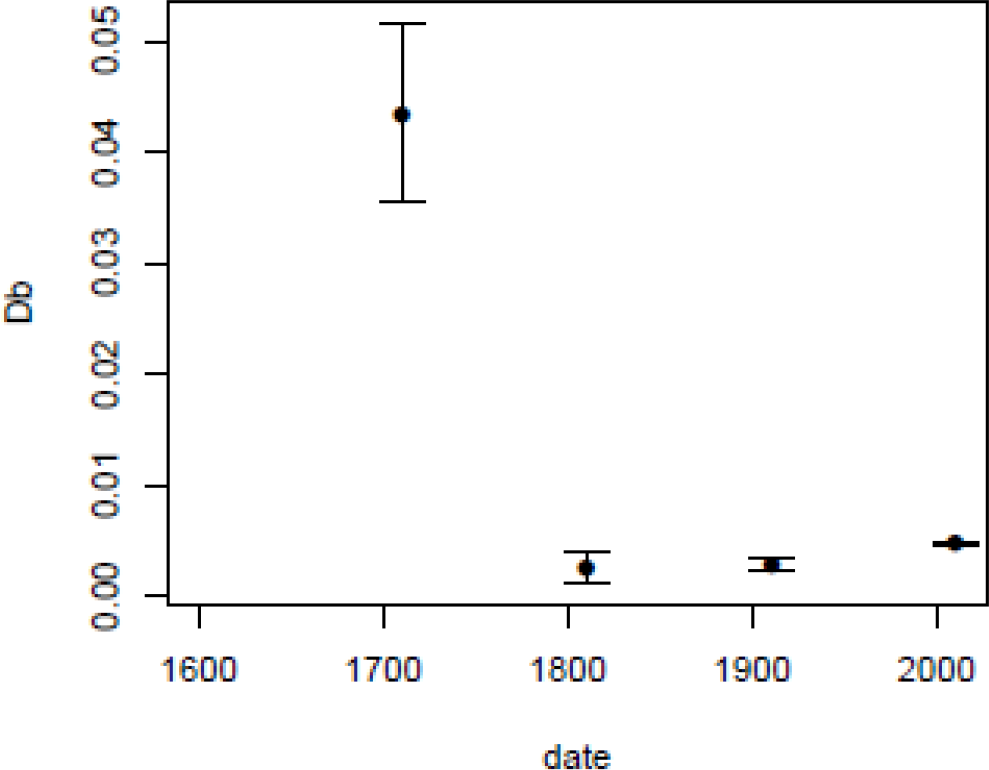
Temporal change in mean pairwise dissimilarity between sites on the Kodiak Archipelago. Error bars represent standard error of the mean.

## Discussion

Studying the evolution of long-lived organisms such as temperate tree species is challenging because of their typically long generation times. This study shows that the concomitant use of tree ring and genetic data can turn the incovenience of these life history characteristics into an opportunity to accurately reconstruct the demographic and genetic history of colonization over five centuries. By applying this combination of methods to several regions and sites within regions at the expansion front of *P. sitchensis*, we were able to describe demographic and neutral genetic patterns of forest establishment at the regional and local scale. We established a link between genetic diversity and spatial colonization patterns by showing that both trends of decreasing diversity and decreasing time since forest establishment towards the expansion front break down at the local scale. This provides insights into the effects of dispersal patterns on neutral evolution during range expansion. It also highlights the importance of relating geographic scale of study to the dispersal abilities of the studied organism when testing for evolutionary trends commonly observed during range expansions. We also established a link between temporal trends in demography and genetic structure of *P. sitchensis* populations over the last five centuries at the expanding range limit: a shift from decreasing to stagnating differentiation between the establishing Kodiak population and the Seward population on the Kenai Peninsula coincides with a marked increase in successful establishment rate on the Kodiak Archipelago, in the 1700s. This suggests that gene flow from continental populations was predominant in early stages of establishment, until local recruitment became the major mechanism of population growth. Most of the allelic richness on the Kodiak Archipelago was acquired during initial stages of population establishment, and the high levels of local gene flow in later stages homogenized the genetic structure of the Kodiak population, buffering founder effects at the local scale and maintaining them at the regional scale.

### Founder and Allee effects

Trees established as early as 1516 were sampled on the archipelago and local recruitment only appears to have become significant in the 1700s, highlighting the existence of a lag of several centuries in local recruitment. Although these results do not provide direct evidence of density-dependent population success, they echo findings of Lesser et al. (2013), who identified Allee effects in early stages of forest establishment in a *Pinus ponderosa* stand through reliance on long-distance seed and pollen dispersal for the first few centuries of population growth. In *P. sitchensis*, wind-borne pollen is likely to be the main vector of genetic material from distant sources, although the abilities of seeds to travel by air and ocean surface currents are not well understood. The shift we observe in differentiation trends in the mid-1700s coincides with the start of a plateau in the allele accumulation curve for the Kodiak-Afognak region. This result can also be related to Roques et al. (2012), who found through models of colonization waves that Allee effects prevent the erosion of genetic diversity along colonization routes: populations at the front accumulate genetic diversity during establishment through reliance on populations behind the expansion front. If a lag in local recruitment through Allee effects is common in populations at the edge of forest tree species, mechanisms described in Roques et al. (2012) could partly explain why many studied forest expansions do not show a classic decrease in diversity at the expansion front. In the case of *P. sitchensis*, when populations had a low density of reproductively mature trees they produced a relatively small pollen cloud compared to external sources, until large mature trees became the main producers of pollen locally present. It is however worth noting that although the expansion process on the Kodiak Archipelago did not result in local erosion of allelic richness, expected heterozygosity over polymorphic sites remained low on Kodiak and Afognak Islands. This suggests that although allelic diversity was recovered largely during colonization, many alleles remain at low frequencies in the newly established population. The genotype of early colonizers might therefore have a long-lasting influence in the establishing population.

### Genetic structure at the expansion front

*P. sitchensis* started establishing at most sites on the Kodiak Archipelago before 1700. Evidence for continental sources dominating gene flow prior to the mid-1700s indicates that long-distance dispersal from the mainland played an important role in initial recruitment. Models of colonization through patches from long-distance dispersal show that genetic sectors are expected to arise (Hallatschek & Nelson, 2010). However, we found no evidence for spatial genetic sectors in the current sample. We propose two explanations for this lack of spatial genetic structure. First, it seems that most founders colonized Kodiak and Afognak Islands from just a few source populations that were already somewhat depleted in alleles. Indeed, it seems that the low allelic richness observed on a regional scale on the Kodiak Archipelago could be due to the low levels of allelic richness of source populations at the tip of the Kenai Peninsula: we observe similar levels of allelic richness on Kodiak and Afognak Islands as in Port Chatham, the closest continental population sampled. Low levels of allelic richness in Port Chatham may be explained by lower population sizes due to a fragmented landscape, as the region is highly mountainous and suitable habitat is restricted to narrow valleys between ice-capped mountains and numerous sinuous fjords. As we also observe higher allelic richness on Shuyak Island than in Port Chatham, multiple source populations might contribute to the maintenance of genetic diversity in regions at the expansion front. The second factor likely to explain the absence of genetic sectors is high gene flow within the Kodiak Archipelago during the last 250 years. We found a decrease in F_ST_ among sites at the expansion front between 1700 and subsequent centuries, suggesting that initial spatial patterns of genetic structure from founding individuals did not persist or develop further due to high levels of subsequent gene flow across the archipelago.

### Demographic estimates of establishment times: power and limitations

Estimates of establishment time, canopy age and time of canopy closure all detected a clear demographic signal of population establishment on Kodiak Island, and a less clear signal for Afognak Island. As these estimates were also calculated for the Seward population, known to be several thousand years old, the power and limitations of these estimates can be established. Mean canopy age is only informative for about 300 years, and estimated time of canopy closure confirmed the recent nature of forests on Kodiak Island and - to a lesser extent - Afognak Island. The complementarity of these two estimates is best illustrated with the Afognak Island site A5: this site is similar to Seward sites in mean canopy age (Figure S1) but juvenile growth rings show a signal of increasing canopy density throughout the 19^th^ century, a pattern not observed in Seward (Figure 2). Although time of canopy closure is useful for inferring the absence of a closed canopy when the first trees established, the temporal change in juvenile radial growth couldn’t be modeled by a logistic curve, or was better fitted by a linear model. This can be due to the initiation of intraspecific competition suppressing growth being too recent, or to spatial heterogeneity within stands. In addition, we found little variability between sites in estimates of time of canopy closure. The parameter *h* from the logistic curve modelling may not be the most informative measure of establishment time. Visually inspecting juvenile ring width over time (Figure 2) might provide more information about stand establishment than extracting a single value from temporal ring width profiles.

### Implications for long-lived wind-pollinated species

Our results suggest that the evolutionary potential of wind-pollinated tree species is more likely to be limited by slow demographic growth than by slow accumulation of genetic diversity. We showed that in spite of a slow initial population growth, allelic richness recovered during this period up to levels comparable to nearby source populations. We also showed that geographic barriers to gene flow are weak despite the studied population being isolated from the continent by a 70 km-wide ocean strait. The demographic lag observed in this and other studies of tree populations suggests an Allee effect, whereby the reproductive ability of establishing trees is limited firstly by a long phase of juvenile growth, and secondly by a higher dependence on foreign pollen fertilizing local mature trees when population densities are low. This potential Allee effect could keep a colonizing tree population in a vulnerable state and contribute to the migration lag of tree species tracking their suitable niche space, especially in the context of rapid anthropogenic climate change. However, such a lag is also likely to be at the origin of an efficient recovery of genetic diversity after a founding event, as several studies have shown a predominance of pollen of foreign origin during early population establishment (Hampe et al., 2013; Lesser et al., 2013; Pluess, 2011). In both cases, management through planting trees from diverse carefully selected provenances could accelerate successful establishment and adaptation in populations of conservation concern (Aitken & Whitlock, 2013). In general, understanding demographic processes during range shifts and their effect on evolutionary potential is necessary as climate change is shifting species’ suitable niches nearly everywhere on the planet. We have shown that dispersal potential and temporal patterns of population growth are important factors influencing population expansion and adaptation, but the effects of other mechanisms such as hybridization, competition, and natural selection also need to be assessed in order to predict or help species movements in response to a rapidly changing world.

## Acknowledgements

This project was funded by a NSERC Discovery Grant to S.N. Aitken and a Strategic Recruitment Fellowship to JS Elleouet from the Faculty of Forestry, University of British Columbia. We thank Christine Chourmouzis, Bert Terhart, Ian Maclachlan, Vincent Hanlon and Jon Degner for valuable help in the field, and Pia Smets for her help in pre-field preparations. Keith Coulter provided generous and indispensable logistic support for collections on Afognak Island. Stacy Studebaker, Bill Pyle, Karl Potts, Ed Berg and Tash Shaheed willingly shared their knowledge about Alaskan forests to help identify sampling sites. Many thanks to Kristin Nurkowski for sharing and helping to refine needle extraction protocols, and Vincent Hanlon for co-developing bark extraction protocols. Lori Daniels and Rafael Chavardès granted access to equipment and patient expertise in dendrochronology methods. Elissa Sweeney-Bergen counted tree rings without counting her hours. Sam Yeaman and Jon Degner provided large bioinformatic support. Thanks to Michael Whitlock, Rafael Cândido Ribeiro and Amy Angert for their comments and suggestions on this manuscript.

## Data accessibility

Tree ring series will be made available on the NOAA website. Genetic data will be deposited on Genbank (raw sequences) and Dryad (SNP dataset) upon acceptance of the manuscript.

## Author contributions

J.S.E and S.N.A designed the study. J.S.E conducted analyses and J.S.E wrote the manuscript with suggestions and modifications from S.N.A.

## Supporting information

Additional supporting information including methods and figures can be found online.

